# Nuclear stiffness decreases with disruption of the extracellular matrix in living tissues

**DOI:** 10.1101/2020.08.28.273052

**Authors:** Kaitlin P. McCreery, Xin Xu, Adrienne K. Scott, Apresio K. Fajrial, Sarah Calve, Xiaoyun Ding, Corey P. Neu

**Author notes:** These authors contributed equally to this work (Kaitlin P. McCreery, Xin Xu).

## Abstract

Reciprocal interactions between the cell nucleus and the extracellular matrix lead to macroscale tissue phenotype changes. The extracellular environment is physically linked to the nuclear envelope and provides cues to maintain nuclear structure and cellular homeostasis regulated in part by mechanotransduction mechanisms. However, little is known about how structure and properties of the extracellular matrix in living tissues impacts nuclear mechanics, and current experimental challenges limit the ability to detect and directly measure nuclear mechanics while cells are within the native tissue environment. Here, we hypothesized that enzymatic disruption of the tissue matrix results in a softer tissue, affecting the stiffness of embedded cell and nuclear structures. We aimed to directly measure nuclear mechanics without perturbing the native tissue structure to better understand nuclear interplay with the cell and tissue microenvironments. To accomplish this, we expanded an atomic force microscopy needle-tip probe technique that probes nuclear stiffness in cultured cells to measure the nuclear envelope and cell membrane stiffness within native tissue. We validated this technique by imaging needle penetration and subsequent repair of the plasma and nuclear membranes of HeLa cells stably expressing the membrane repair protein CHMP4B-GFP. In the native tissue environment *ex vivo*, we found that while enzymatic degradation of viable cartilage tissues with collagenase 3 (MMP-13) and aggrecanase-1 (ADAMTS-4) decreased tissue matrix stiffness, cell and nuclear membrane stiffness is also decreased. Finally, we demonstrated the capability for cell and nucleus elastography using the AFM needle-tip technique. These results demonstrate disruption of the native tissue environment that propagates to the plasma membrane and interior nuclear envelope structures of viable cells.

## INTRODUCTION

Understanding biological processes that coordinate tissue development, pathology, and regeneration requires the study of living systems on multiple scales, from gene regulation within the nucleus of a single cell to the synchronization of cell and tissue networks. Changes in cell gene expression lead to alterations in biochemical signaling that regulate cell communication, timing of cellular activities, and ultimately, tissue structure. At the same time, the extracellular matrix (ECM) dictates another layer of cellular regulation. Biochemical cues, physical forces and changes in stiffness of the ECM microenvironment guide cell migration, proliferation, differentiation, and changes in gene expression ^1–5^. Therefore, the response of the cell to the native microenvironment is an emergent behavior that connects micro- and macro-scale tissue architecture with cell distribution, proliferation, and differentiation.

Pathological tissue processes occur across multiple biological length scales, impacting the tissue matrix and the cell nucleus alike. Cells are physically linked to their local matrix via focal adhesions, allowing cells to respond the physical environment ^6^. Within the cell, the cytoskeleton is connected to the nucleus through the LINC (Linker of Nucleoskeleton and Cytoskeleton) complex. As a result, variations in ECM mechanics are propagated to the cell nucleus affecting cellular processes ranging from protein conformation, localization of transcription factors, and chromosome organization ^7–9^. The cell nucleus is a large, stiff organelle that regulates homeostasis and cell phenotype partly through mechanotransduction mechanisms ^10^. Within the nuclear envelope, a mesh of linked lamin proteins interact with cytoskeletal and nuclear actin and act as a shock absorber to maintain nuclear architecture when the cell environment changes^11,12^. Chromatin and lamin proteins are main contributors to the mechanical properties of the nucleus, and additionally have both a role in gene regulation and mechanotransduction mechanisms ^13^. Chromatin, which fills most of the volume of the nucleus, is more or less viscoelastic depending on the ratio of heterochromatin to euchromatin, which is controlled by gene regulating proteins ^9,14^. Therefore, studying the mechanical properties of the nucleus can provide insight into gene regulation that could be critical for characterizing changes in cell phenotype or cell pathology. Although nuclear mechanics has been explored *in vitro* ^15,16^, the complex interplay between matrix, cell, and nuclear stiffness in developing and pathological tissue states is not holistically understood, largely due to experimental limitations.

To elucidate multiscale tissue function and pathology, the mechanics of *ex vivo* tissues has been investigated using a variety of techniques ^17^. Stress relationships within tissues can be determined by injecting fluorescent markers, such as liquid droplets, into the matrix space and tracking their displacement and deformation when defined forces are applied ^18^. Tissue properties can also be inferred using embedded magnetic beads to track displacement when a magnetic field is applied ^19^. On a smaller physical scale, spatiotemporal stiffness of chromatin within the cell nucleus can be extracted with image-based nuclear elastography ^20^, though these approaches have not yet resolved the interplay of nuclear, cellular, and matrix properties within viable tissues. Rather than extracting relative mechanical relationships such as relative stiffness and tissue viscosity, direct mechanical contact manipulation of cells and tissues use controlled forces to push or pull on a sample facilitating the direct extraction of mechanical properties. This approach employs tools such as indenters or micropipette aspiration to extract cell stiffness ^21,22^. These methods have been independently applied to sustained viable tissues and cells in culture, but have not yet been developed to distinguish the contributions of cells, nuclei, local matrix, and ECM components in a single tissue sample. Atomic force microscopy (AFM) has been applied to biological systems in recent decades, and with basic techniques, can extract mechanical properties of local matrix and cell stiffness directly in viable tissues ^23,24^. AFM techniques have been modified to measure nuclear mechanics in living cells ^22^, but not yet while the cells are still embedded in native matrix to investigate the multiscale pathology of tissue disease states.

Here, we report a method to investigate the impact of ECM degrading enzymes on nuclear mechanics while cells are embedded in their native tissue environment. We combine atomic force microscopy and confocal fluorescent microscopy to facilitate cell and nuclear force-spectroscopy, allowing us to directly probe specific regions of the same viable tissue sample, including the cell nucleus, cell membrane, and the corresponding extracellular space. We applied these methods to measure *ex vivo* cartilage tissue sections after treating with matrix-degrading enzymes MMP-13 and ADAMTS-4 to investigate the degradation of the cartilage matrix on nuclear stiffness. The experimental design allowed us to take stiffness measurements of nuclear, cell, and local extracellular matrix without disrupting nuclear/cell/matrix interactions. Since the nucleus is a mechanosensitive organelle that can remodel itself to maintain homeostasis and protect intranuclear machinery ^25^, we hypothesized that nuclear stiffness would decrease to match biomechanical disruption of the extracellular environment caused by enzymatic degradation of the cartilage ECM. Our results suggest that enzymatic degradation of cartilage tissues decreases matrix stiffness that is subsequently propagated to the nuclear envelope, leading to significant changes in nuclear membrane stiffness. These results demonstrate direct measurements of plasma and nuclear membrane stiffness while cells remain in their native tissue environment.

## RESULTS AND DISCUSSION

The nucleus bears mechanical loads from the surrounding ECM and responds to tissue degradation. Thus, it is important to observe and measure the cell and its nucleus in the native tissue environment, as nuclear and cellular mechanics are profoundly influenced by tissue processing ^23^. While AFM needle-tip techniques have been used to directly measure nuclear stiffness of isolated nuclei and isolated cells ^22^, we report a subsequent advanced technique to directly measure nuclear membrane stiffness while maintaining cell-matrix interactions. The question addressed by the present study is whether biochemical treatment and biomechanical disruption of tissues is transmitted to the cell and nucleus, and how the responses of the nucleus to the ECM may be captured using the AFM needle-tip technique. These experiments show that enzymatic treatment of cartilage tissue explants causes a softening of nuclei within embedded cells, and that the local extracellular matrix, cell membrane, nuclear membrane stiffnesses may be distinguished using these methods.

This investigation employs an AFM needle-tip probe to directly measure cell and nuclear membrane stiffness while cells are viable and sustained in their native tissue environment (**Figure 1**). The AFM needle-tip probe is capable of measurements with high physical resolution because the tip interacts with only the targeted biological feature spanning the tip contact point, while larger probes will yield properties that average out over the indented area (**Figure 1A**). We use fluorescence microscopy to visualize cell and nuclear structures in order to align the needle tip over a cell and its nucleus or the local ECM region within 10 μm of the cell body. We characterized the approach and retraction of the AFM needle-tip to/from the cell and nuclear structures into five distinct stages (numbered in **Figure 1B**). These stages can be matched to the acquired AFM data to interpret material behavior upon indentation of the cell (corresponding numbers in **Figure 1C**). As the AFM needle approaches the cell in (1), the curve segment is flat because the AFM needle has not yet contacted the cell. During (2), the needle contacts the cell membrane and deforms it until puncture, causing the observed force relaxation. As the cantilever is lowered further, the needle loads the nucleus and punctures the nuclear envelope denoted by step (3). Segments at (2) and (3) are used to interpret the stiffness of the plasma membrane and nuclear envelope simply by calculating the slope of the peak before relaxation. The final increasing slope at the end of the approach curve (4) can be attributed to the interaction between the needle tip and intranuclear components, such as the nucleoskeleton and chromatin, as they deform upon indentation. Once the setpoint voltage is reached, which is preset by the user corresponding to the maximum force imposed on the substrate by the AFM cantilever, the cantilever is triggered to retract from the nuclear and cell structures (5).

**Figure 1.**
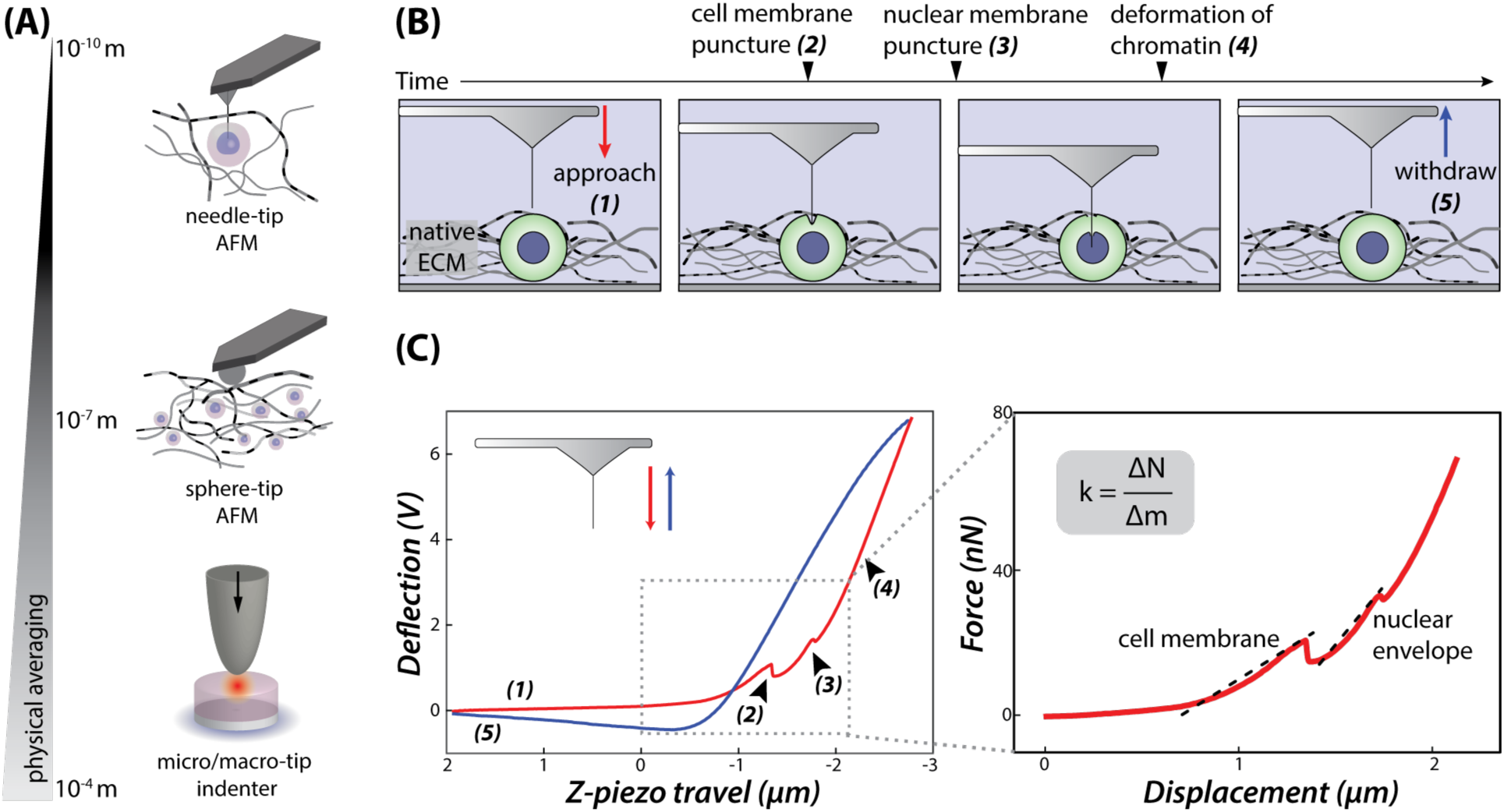
Probing the cell nucleus of viable cells in tissues with needle-tip AFM probe. (A) Varied probe geometry and sizes may be used to contact and probe the mechanics of biological samples. Larger indenters, on the order of 10^−4^ meters (bottom), will probe samples and average biological features most over a sample area. Sphere-tip probes, on the order of 10^−6^ meters, can probe the surface of biological specimens. A pyramidal tip sharpened to a needle on the order of 10^−9^ meters is the finest physical probe size, able to puncture biological membranes to investigate cell and nuclear mechanics. (B) Schematic of experimental methods to probe cell and matrix stiffness while cells are still embedded in native tissue, illustrating (1) the approach, (2) the needle puncturing the cell membrane, (3) the needle puncturing the nuclear membrane, (4) the deformation of chromatin and intranuclear components, and (5) retraction of the needle from the nucleus and cell bodies. (C) Experimental AFM data collected using needle-tip AFM on a chondrocyte embedded in its natural cartilage tissue matrix. Two distinct force relaxations in the approach curve correspond to cell membrane puncture and nuclear membrane puncture, respectively. By fitting the incline of the force-displacement before relaxation to a linear model that is independent of probe interactions with the sample, feature stiffness can be extracted.

To determine the stiffness of the cell membrane and nuclear envelope using experimental AFM puncture curves, one must consider the material response of the biological membrane both to external ECM forces and probe contact. Generally, AFM data is used to extract material properties of biological materials by fitting the AFM indentation curve to the Hertz theory of elasticity to extract Young’s modulus ^26^. However, the native ECM exerts an isotropic pre-stress tension through cell-matrix attachments which oppose the applied force from the AFM probe during membrane deformation, limiting the applicability of typical force-indentation models for elasticity ^27^. Furthermore, an accurate derivation of elastic modulus for a material requires a careful estimation of the AFM tip geometry ^28^. Unfortunately, precise tip radius cannot be routinely characterized for the needle-tip probes as they are fabricated from a spontaneous growth of Ag_2_Ga crystals normal to the cantilever surface, resulting in an approximate tip radius of 20 nm — 100 nm ^29^. At the same time, when indenting a cell membrane with a sharp AFM tip, the bending resistance of the membrane dominates the material response ^30^. Taking this contextual evidence into account, we report stiffness values of the cell membrane and nuclear envelope by fitting the force-displacement data before relaxation of a corresponding needle puncture to a linear model (**Figure 1C**). These data are dependent on the applied force from the AFM probe and deformation only, reflecting an effective bending stiffness of the biological membrane structures when indented by a needle probe.

To validate needle penetration into the cell and nuclear structures, we mounted the AFM system onto an inverted laser scanning confocal microscope to observe both the characteristic force spectroscopy curve and simultaneously image fluorescence of isolated cells (**Figure 2**). For these experiments, we used a HeLa cell line stably expressing a charged multivesicular body protein 4B (CHMP4B-GFP) to distinguish membrane deformation with membrane puncture. Cells require CHMP4B to repair damaged plasma and nuclear membranes ^31,32^. When tagged with fluorescence, confocal microscopy can be used to visualize the localization of CHMP4B to nuclear envelope and plasma membrane after physical disruption ^33^. By tracking localization of CHMP4B, we show that the integrity of the plasma membrane is repaired within 10 minutes of puncture with the AFM needle-tip probe. To visualize the probe during the membrane puncture, we coated the AFM needle-tip probe with fluorescently labeled fibronectin (**Figure 2B**). A constant force was maintained at the setpoint when probing HeLa CHMP4B-GFP to capture *z*-stack and planar images of the needle positioned within the cell nucleus (**Figure 2**).

**Figure 2.**
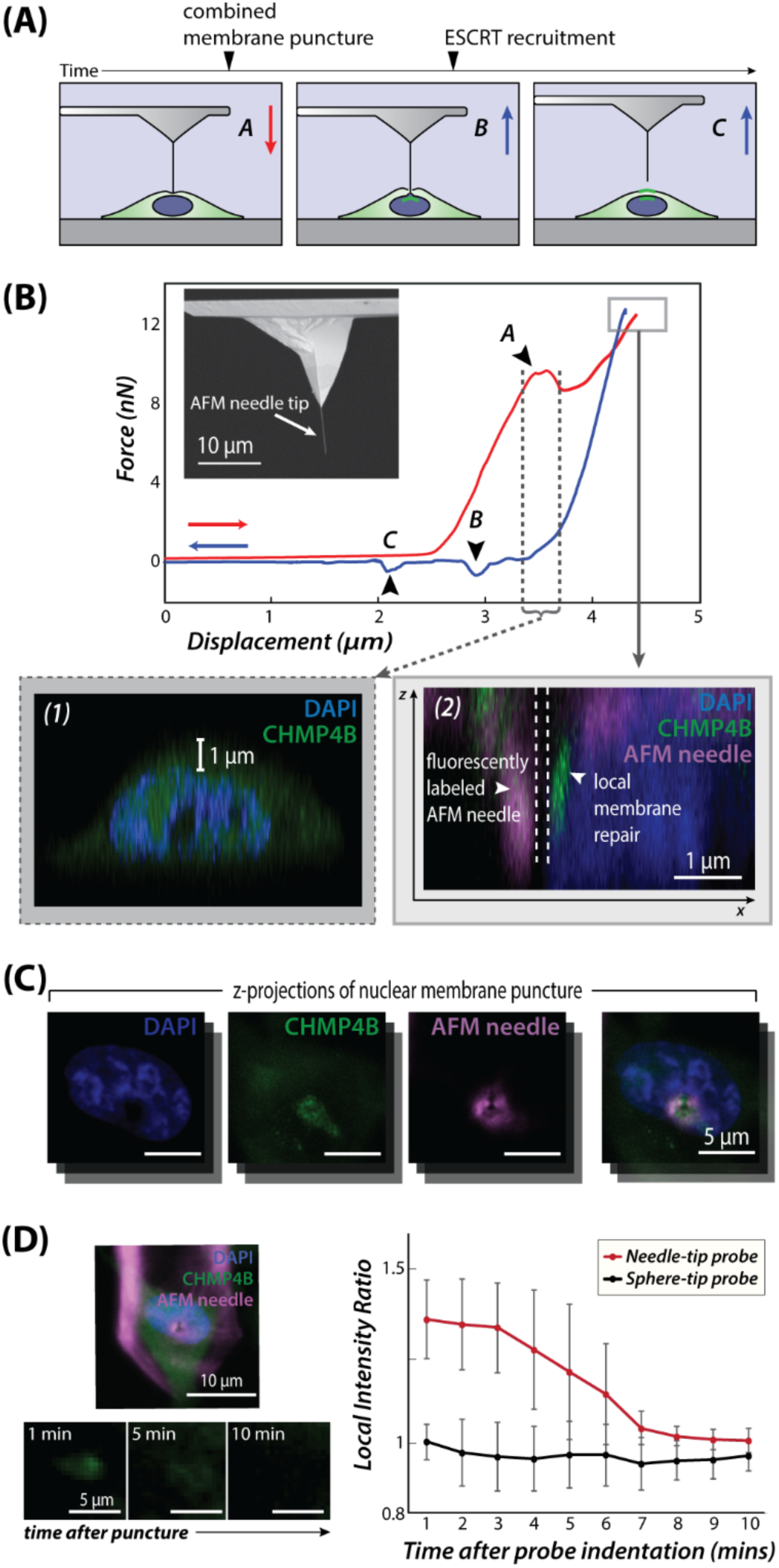
Visualization and validation of dual membrane puncture via CHMP4B-GFP recruitment and fluorescence microscopy. (A) Schematic of probing isolated HeLa cells stably expressing CHMP4B-GFP with a needle-tip AFM probe. (B) SEM image of AFM needle-tip probe, indicating the location of the needle. Corresponding experimental data for force-spectroscopy measurements indicating force-relaxation regions during A: combined membrane puncture, B: retraction of probe from nuclear membrane, and C: retraction of probe from plasma membrane. A fluorescent Z-stack reconstruction before (B1) needle puncture reveals the small (∼1 μm) distance between plasma and nuclear membranes, matching observations of the force-relaxation curve. Indirect visualization of the AFM needle-tip (pink) and CHMP4B recruitment at the puncture site (green) within the nucleus (blue) during needle puncture (B2). (C) Projected *z*-stack image spanning 10 μm obtained during cell puncture allowing simultaneous visualization of cell nuclei (blue), CHMP4B recruitment (green), and indirect needle probe staining (pink). (D) Relative fluorescence intensity, defined as the ratio of GFP intensity at the puncture site relative to a site 10 pixels above in the *xy*-plane, at the location of the force-spectroscopy grid (4×4 indentations, 5×5 μm) of HeLa CHMP4B-eGFP cells (n=10) probed with a needle-tip (red) or sphere-tip (black) probe.

During HeLa cell puncture, force-displacement curves typically demonstrated a single force peak before relaxation upon probe approach, and two distinct relaxation segments when retracted (**Figure 2B**), in contrast with chondrocyte membrane puncture depicted in **Figure 1**. This single peak phenomenon was most likely observed because of the small distance (< 1 μm) between the cell and nuclear membranes typical of a HeLa cell, causing the membrane to deform and evenly contact the nuclear envelope before puncturing both (**Figure 2A**). A combined membrane stiffness, effectively a series of mechanical resistance, can be directly calculated from the approach curve. In contrast, the cell membrane and the nuclear membrane can be distinguished with AFM measurements of chondrocyte cells because there is a large space between the nucleus and the plasma membrane (**Figure 1B**). This finding presents a limitation of the AFM needle-tip technique because the nuclear membrane stiffness may not be resolved in cells that spread on culture plates or have plasma membranes that are easily deformable. Upon retracting the needle probe, two relaxation curves correspond to the exit of the needle tip from the nuclear and cellular membranes respectively, confirming that both structures were indeed punctured with the needle.

Fluorescence imaging reveals that CHMP4B recruitment occurs at the site of the needle within the cell nucleus following nuclear membrane puncture (**Figure 2C**), confirming previous studies that ESCRT-III machinery is involved in both nuclear and plasma membrane repair when physically disrupted ^32,33^ (**Figure 2D**). While the needle tip is only ∼20 – 100 nm in diameter, multiple punctures within a small area (4×4 points within 5×5 μm^2^) reveal more prominent CHMP4B-GFP foci at the puncture site for observation. The CHMP4B activity is visible at the wound site less than 1 minute after the spectroscopy scan. After 5 minutes of membrane repair, the fluorescence intensity decreases, but is still visible. At 10 minutes post-disruption, the CHMP4B-GFP intensity returns to baseline levels as the plasma membrane repairs are complete. For comparison, a sphere-tip probe used to indent cell membrane structures demonstrated little change in fluorescence intensity at the site of measurements, confirming that the cellular membrane is not disrupted by a rounded tip and only requires repair following interaction with the needle-tip probe. These results clearly show cell functionality and viability during and immediately after needle penetration. While long-term effects and cell viability after needle puncture were not assessed in this study, existing evidence suggests that cells are recoverable and remain viable after AFM needle tip puncture ^22^. Taken together, these procedures indicate that live cells and their nuclei may be probed directly by the AFM needle-tip technique.

Next, we investigated the impacts of biochemical degradation of bovine cartilage tissue on multiscale (ECM, cell, and nuclear) tissue stiffness when treated over time with one of two cartilage degrading enzymes using the AFM needle tip technique combined with fluorescence microscopy. Articular cartilage is composed of chondrocytes that secrete and regulate turnover of matrix proteins that maintain tissue homeostasis. Cartilage disease states, such as osteoarthritis (OA), cause chemical and mechanical degradation in joint tissues attributed to altered interactions between the chondrocyte and its local environment. One of the early biochemical changes to cartilage following traumatic injury or degenerative joint diseases is the breakdown of aggrecan and type II collagen, two major structural components of articular cartilage ^34^. Type II collagen forms a fibril network and provides tensile strength to cartilage, while aggrecan is a structural macromolecule involved in fluid retention to provide resistance to compression ^35^. Two enzyme families associated with degrading the majority of the collagen and aggrecan in human arthritic cartilage include the enzymes Matrix MetalloProteinases (MMPs) and A Disintegrin And Metalloprotease with Thrombospondin motifS (ADAMTS) ^34^. ADAMTS-4 is a principle aggrecanase in murine, bovine, and human articular cartilage responsible for the degradation of aggrecan and other proteoglycans in joint disease ^36^. ADAMTS-4 expression is upregulated in OA chondrocytes and is responsible for pathological cleavage of aggrecan that results in the accumulation of large proteins in the joint ^37,38^. Meanwhile, MMP-13 cleaves interstitial fibrillar collagens among other ECM molecules, but primarily cleaves type II collagen in later stages of OA ^37,39^. MMP-13 is also responsible for cleaving other structural molecules in cartilage including type IV, X, and XIV collagens, fibronectin ^40^, fibrillin-1 ^41^, perlecan ^42^, and aggrecan ^43^. While the upregulation of these proteases is linked to biochemical and biomechanical disruption in the joint resulting from disintegration of key matrix components, the interplay between matrix degradation and matrix-producing chondrocytes is only beginning to be explored.

While these methods may be used in the context of other tissues, cartilage is a suitable choice due to the ease of *ex vivo* culturing and ability to target the degradation with matrix-specific enzymes. It is the relatively avascular and aneural properties of cartilage that makes it a suitable tissue choice for an *ex vivo* culturing system to probe factors related to cartilage degeneration and regeneration ^44^. Tissue sections parallel to the cartilage surface were sliced using a vibratome to expose rounded, middle-zone chondrocytes. We located live chondrocytes and the chondrocyte nuclei by staining cells with vital Calcein AM and DAPI. After probing cell structures with the AFM, we measured nearby ECM stiffness outside of the chondron, hereon referred to as the local extracellular matrix to the cell. On all physical scales measured, there was no significant effect of culture time on the stiffness of untreated cultured cartilage tissues (p < 0.05). Thus, change in stiffness is the result of matrix degradation, and not an artefact of culturing tissues *ex vivo*. Additionally, the matrix-targeting enzymes used in these studies are not considered to impact cell and nuclear behaviors directly. In healthy cartilage, chondrocytes constitutively express and secrete MMP-13, which acts to cleave a range of type II collagen peptides for tissue maintenance and remodeling, and is then rapidly endocytosed and degraded by chondrocytes ^45^. ADAMTS-4 is usually present during inflammation, but is then cleared from the extracellular space via endocytosis ^46^. Because chondrocytes regulate extracellular enzymes, and enzyme activity occurs in the extracellular space, cell and nuclear stiffness values reported here are likely reflective of the ECM mechanics and not a direct result of cellular exposure to MMP-13 and ADAMTS-4. However, the impact of higher enzyme doses on cell and nuclear mechanics may be the topic of future studies.

We found that disruption of the cartilage ECM by ADAMTS-4 results in decreased stiffness of cartilage, the cell membrane and nuclear envelope of embedded chondrocytes (**Figure 3**). After 7 days of co-culturing cartilage tissue explanted from 3 bovine animals with ADAMTS-4, the bulk stiffness of cartilage plugs is significantly decreased by day 4 compared to the control, which confirms the efficiency of this aggrecanase to alter the bulk biomechanical integrity of cartilage tissue (**Figure 3A**). After 7 days of treatment with ADAMTS-4, probing the nuclear, cellular, and local matrix structures with needle-tip AFM further revealed changes in stiffness of the matrix and the cell. Not only did the local matrix stiffness significantly decrease with ADAMTS-4 on this scale (p < 0.05), but the cell membrane stiffness is significantly reduced by 51.72% compared to the control (p < 0.05). This could be explained by the close association of aggrecan and proteoglycans to the cell membrane. Aggrecan is indirectly bound to the cell surface via hyaluronan, so its disruption would be detrimental to fluid retention within the chondron just outside of the plasma membrane ^47^. Within these cells, we found that ADAMTS-4 treatment significantly reduces nuclear envelope stiffness by 44.1% compared to the control (p < 0.05) (**Figure 3C**). This provides evidence that ADAMTS-4 disrupts the cartilage matrix, and this disruption is transmitted to the cell nucleus.

**Figure 3.**
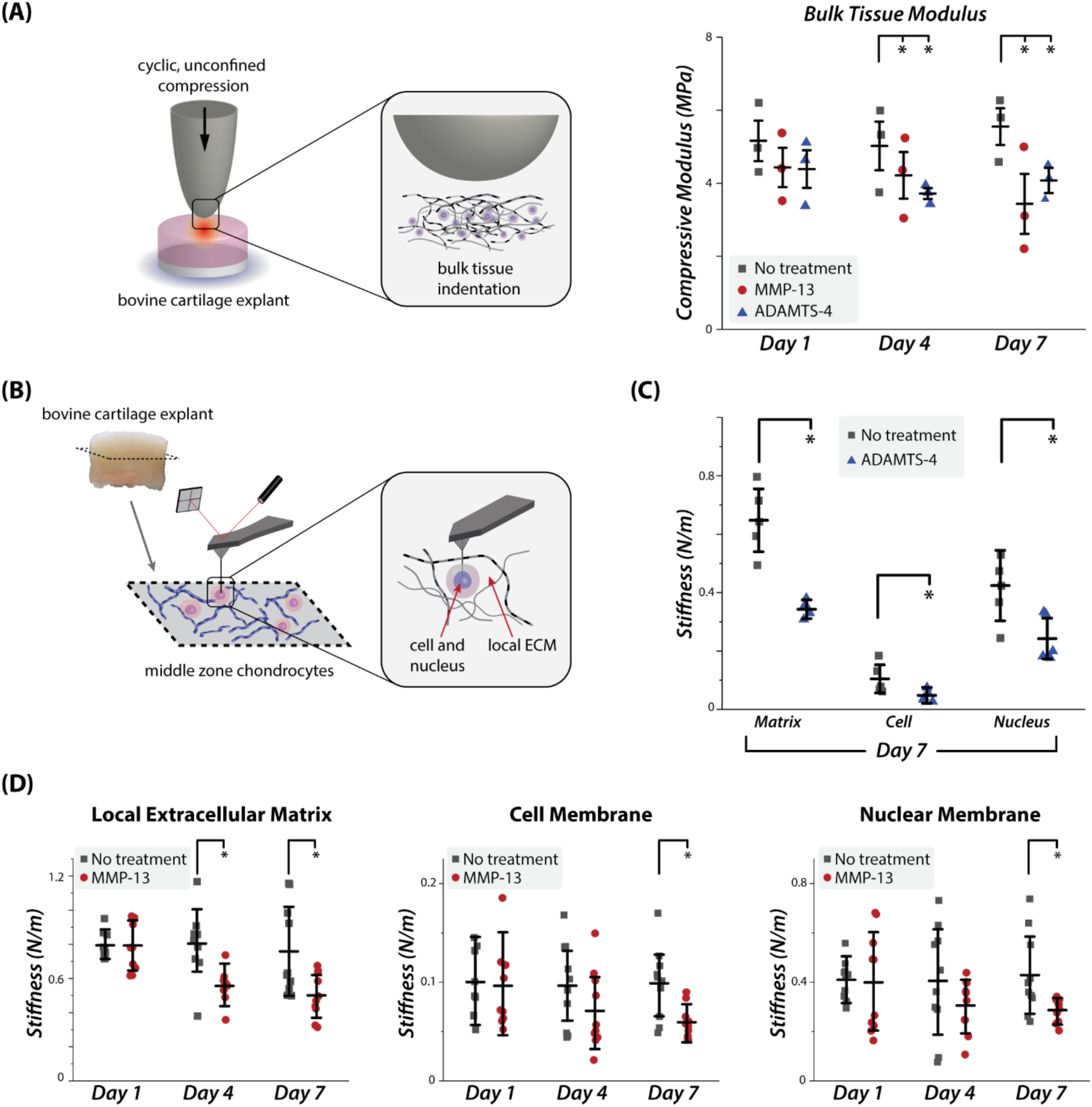
Multiscale mechanical properties of cartilage tissue, embedded cells, and their nuclei after *ex vivo* tissue treated over time with matrix degrading enzymes MMP-13 and ADAMTS-4. (A) Cyclic, unconfined compression using a 1mm indenter determines averaged stiffness of cartilage tissue. Compressive modulus of cartilage tissue (*n* = 3 animals) treated with MMP-13 and ADAMTS-4 after 1, 4, or 7 days incubation. (B) After slicing cartilage perpendicular to the articular surface, AFM needle-tip probe facilitates measurements of chondrocyte cell and nuclear membrane stiffness while cells are still embedded in native ECM. (C) Local matrix outside the chondron, chondrocyte cell, and nuclear membrane stiffness (n = 5 cells) after treatment of cartilage tissue with ADAMTS-4 for 7 days. (D) Local ECM, cell membrane, and nuclear membrane stiffness after tissue treatment with MMP-13 for 1, 4, or 7 days (*n* = 10 cells). *p < 0.05 for all pairwise comparisons and values are reported as mean ± standard error of the mean.

Additionally, biochemical disruption of the matrix by MMP-13 reduces the stiffness of the cartilage tissue, local matrix, cell membrane, and nuclear envelope (**Figure 3D**). MMP-13 is the major collagenase in OA cartilage, typically active in late stages of OA degeneration ^37^. After co-culturing cartilage tissue explants with MMP-13 for 1, 4, or 7 days, the bulk tissue stiffness was significantly reduced by day 4 (p < 0.05) though not significantly reduced between 4 and 7 days (p > 0.05), similar to the results we observed after treatment with ADAMTS-4 (**Figure 3A**). Probing the local extracellular matrix stiffness with needle-tip AFM showed significant decrease by day 4 of incubation (p < 0.05), but the difference plateaus by day 7 and is not significant between days 4 and 7, confirming the time course findings in bulk testing studies. Finally, cell membrane stiffness was reduced by 40.01% compared to the control by day 7, and the nuclear membrane stiffness was reduced by 32.56% (p < 0.05). However, we did not find significant evidence of reduced stiffness in either of these structures before day 7 (p > 0.05). Therefore, as MMP-13 disrupts ECM molecules in the matrix and reduces its foundational structure, one cellular response is reduction in nuclear stiffness.

Each enzyme treatment causes a significant decrease in nuclear stiffness by day 7, but their discrepancies provide insight into the propagation of matrix signals to the cell nucleus. Compared to each respective control, the mean nuclear stiffness among cells in tissues treated with MMP-13 decreased by 32.56%, while the mean nuclear stiffness in ADAMTS-4-treated tissues decreased by 44.13% (p < 0.05). These discrepancies may be partially explained by the impact of disrupting different functional components of cartilage. MMP-13 preferentially targets type II collagen as a primary structural molecule of articular cartilage that imparts a tensile strength to balance the natural swelling behavior of the tissue. MMP-13 cleavage of type-II collagen may additionally perturb ECM-cell interactions by disrupting the collagen-mediated cell attachment to the matrix via integrin receptors and non-integrin collagen receptors ^48,49^. Meanwhile, ADAMTS-4 degrades aggrecan, a matrix molecule that contributes to cartilage swelling, and additionally the influence of aggrecan bound to hyaluronan, which tethers the pericellular matrix to the cell surface ^47^. Moreover, disruption of aggrecan in close association with the chondrocyte membrane, as well as proteoglycans in the pericellular matrix, are likely to have a severe impact on cell matrix stiffness as large GAGs accumulate and fluids cannot be efficiently retained surrounding the cell body. As a result, disruption of the cartilage matrix destabilizes the multiscale cartilage structure and the transfer of intrinsic forces within the tissue, which may then also be transmitted to the cell surface. Disruption of the links between the extracellular matrix to the cell nucleus is a major contributor to the pathology of disease by triggering chromatin rearrangement, ultimately having a major effect on gene expression ^50^. These findings represent only the beginning of nucleus-matrix interactions that may be investigated with direct probing of nuclear stiffness in viable tissues using the AFM needle-tip technique.

Unlike isolated cells sustained in culture, cells embedded in their native tissue matrix are subject to prestress from attachments to the surrounding ECM which influences both plasma and nuclear membrane stiffness. The biomechanics of needle penetration is optimized by pre-tensioning the biological membrane with externally applied forces ^51^. Cells embedded in their native tissue environment are subject to prestress of the plasma membrane, which is linked to the ECM by cell surface receptors ^52^. The nuclear envelope is analogously prestressed by the cytoskeleton through the LINC complex, which physically tethers the nucleus to cytoskeletal F-actin ^53^. As a result, both the cell membrane and nuclear envelope are subject to tensional and compressional stress critical to maintaining nuclear function, and balancing these opposing forces directly impacts cell and nuclear deformability ^54^. Cells compensate for reduced prestress through cytoskeletal remodeling, which results in changes of prestress applied to the nucleus and has the potential to create vulnerability in the nuclear envelope ^55^. In this study, we altered prestress applied to the cell exterior by treating cartilage tissues with matrix-degrading enzymes, which consequently impacted cell and matrix penetration mechanics. By maintaining the interconnectivity between the nucleus and the cytoskeleton, and the cytoskeleton and ECM, our results indicate that prestress disruption is propagated between these regions. While other tissue types were not featured in this study, cell stiffness may be different in other tissues due to the existing prestress that is more or less isotropic or uniaxial depending on the matrix structure. Thus, our conception of nuclear stiffness should not be limited to quantification of nuclear mechanics alone or nuclear mechanics in living cells, but should encompass interactions of the nuclear envelope with the extracellular prestress among tissue types.

Finally, a preliminary study of cell and nucleus elastography was conducted on isolated primary chondrocytes using a needle-tip AFM probe (**Figure 4**). Stiffness maps of a cell and its nucleus are generated by a force-spectroscopy scan, where each pixel in the membrane and nuclear regions is an independent puncture curve (**Figure 4A**). By combining fluorescent microscopy and the AFM needle-tip technique, a stiffness map of an isolated cell may be aligned and overlaid with microscopy images. These data set the stage for future studies of nuclear elastography of cells within living tissues. To accomplish this, several challenges remain that will be unique to tissue types. For example, articular cartilage cells may be buried under fibrous matrix, which varies in depth across one cell and between cells posing a potential hurdle for AFM scanners equipped with a limited vertical travel distance. This phenomenon impeded our initial attempts to map chondrocyte cell and nuclear stiffness *in situ*. However, direct measurement of nuclear mechanics at this resolution *in vitro* would facilitate the correlation of nuclear stiffness with fluorescent reporters indicative of architecture and composition when combined with simultaneous fluorescent imaging. Furthermore, this method will enable the ability to monitor changes in nuclear mechanics with fluorescent indicators in real-time. With methodical modifications and application, nucleus elastography facilitated by the AFM needle-tip technique may be applied in viable tissues in future studies to investigate the relationship between nuclear functionality and pathological state at higher resolution.

**Figure 4.**
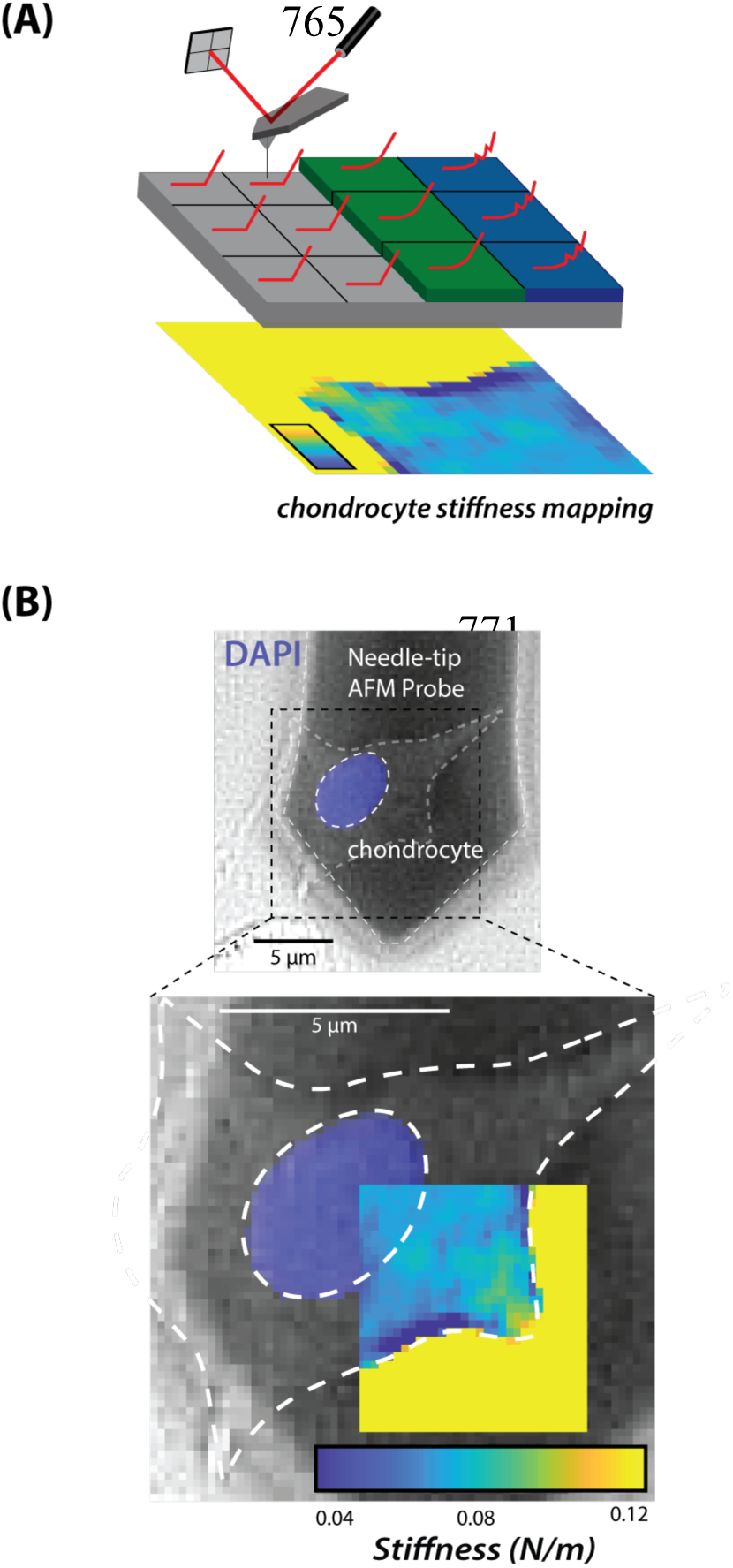
Mapping isolated cell and nuclear membrane stiffness via needle-tip AFM elastography. (A) Force-spectroscopy mapping of isolated chondrocyte cells generates a high-resolution stiffness map via multiple puncture curves of the cell and nuclear membranes and indentation of surrounding regions. (B) Experimental data mapping the stiffness (colored) of a viable isolated chondrocyte (P0) cultured on fibronectin-coated glass, simultaneously imaged via fluorescence microscopy (greyscale, DAPI). In the colored map, blue denotes the softest areas and yellow the stiffest areas.

## CONCLUSION

We demonstrate that needle-tip AFM probes facilitate direct probing of cell and nuclear membrane stiffness among viable cells in their native tissue environment. Force-displacement data are obtained when the AFM needle tip deforms and then punctures the cell membrane and nuclear envelope, which can be achieved in viable tissues when combined with *ex vivo* culture conditions and fluorescence microscopy. The results presented show a significant decrease of nuclear stiffness as a result of biochemical matrix degradation, elucidating cell response mechanisms to the extracellular tissue environment. Disruption of type-II collagen has a significant effect on nuclear stiffness, perhaps due to the loosening of macromolecular matrix proteins and the disruption of key integrin-mediated links between the chondrocyte and its local matrix. Plasma membrane and nuclear stiffness decreases by degradation of aggrecan via ADAMTS-4, likely observed because aggrecan is bound to the cell surface and retains compressive resistance. Overall, the main advantage of the AFM needle-tip technique is that it may be used to gain insight into the mechanisms by which cell nuclei differentially respond to biochemical and biophysical stimuli during tissue development, disease, and regeneration. By studying changes of mechanics of nuclei, we may gain insight into how chromatin rearranges and is regulated in specific mechanical environments. Additionally, our AFM technique is able to probe physiological and pathological changes in tissue environments which influence cell function, nuclear mechanotransduction mechanisms, and the interaction between the extracellular matrix and nuclear mechanics.

## MATERIALS AND METHODS

### Cell and Tissue Explant Culture

All cell and explant tissue cultures described below were sustained in chemically defined Dulbecco’s modified Eagle medium nutrient mixture F12 (DMEM; Life Technologies, Grand Island, NY, USA) and supplemented with 10% fetal bovine serum (Life Technologies, Grand Island, NY, USA), 1% bovine serum albumin (Sigma-Aldrich, St. Louis, MO, USA), 50 ug/mL ascorbate-2-phosphate, and 1% penicillin and streptomycin (Life Technologies, Grand Island, NY,USA) unless stated otherwise.

Cartilage tissues were extracted from bovine stifle (knee) joints from 2-week old calves within 12 hours of slaughter. The joints were opened under aseptic conditions, exposing femoral condyles. Osteochondral plugs were removed from the load-bearing regions of distal femoral condyles with an 8 mm diameter coring reamer ^56,57^. Briefly, the plugs were washed thoroughly with sterile phosphate-buffered saline (PBS; Life Technologies, Grand Island, NY, USA) and incubated in the culture medium at 37 °C and 5% CO_2_ for 24 hours for equilibrium prior to enzyme treatment. For cell extraction, chondrocytes were isolated by digestion with 0.2% collagenase-P (Roche Pharmaceuticals, Nutley, NJ) for 5 hours. Digested tissue was then filtered with a 70µm cell strainer (Fisher Scientific, Hampton, NH, USA). Isolated cells were then washed (3×) with standard supplemented culture media. The resulting chondrocyte cell population was cultured in this modified media at 37 °C with 5% CO_2_ until AFM testing 2 days later.

HeLa cells expressing CHMP4B–GFP were a gift from the Martin Stewart Lab, Max Planck Institute, Dresden, Germany ^58^. Cells were routinely cultured in 75 T flasks at 37 °C with 5% CO_2_ in standard supplemented culture media with an additional supplement of 1 % of geneticin (Sigma-Aldrich, St. Louis, MO, USA).

### Enzyme treatment and tissue preparation of bovine cartilage live tissue

Recombinant human ADAMTS-4 (R&D Systems, Minneapolis, MN, USA) and MMP-13 (R&D Systems, Minneapolis, MN, USA) were diluted to 50 ng/ml in the tissue explant culture medium. Cartilage explant tissues were incubated in the control culture medium and 50 ng/ml ADAMTS-4 at 37 °C and 5% CO_2_ for 7 days; or incubated in the control culture medium and 50 ng/ml MMP-13 at 37 °C and 5% CO_2_ for 1, 4 or 7 days respectively. The plugs were then sliced perpendicular to the articular surface in 30 μm-thick sections by a vibratome (VT-1000S, Leica Microsystems Inc., Germany) in physiological buffer to preserve cell viability. Cartilage sections were washed thoroughly with PBS and pre-stained with Calcein AM and DAPI (Life Technologies, Grand Island, NY, USA) to identify live cells, and then affixed to a coverslip by applying a small drop of cyanoacrylate (Loctite, Westlake, OH, USA) to each end of the tissue sections but not the AFM testing regions as previously described ^23^.

### Bulk Measurement of Cartilage Explants and Data Analysis

Cartilage plugs were tested in unconfined compression using a bench-top mechanical testing system (ElectroForce^®^ 5500 Test Instrument, TA Instruments, Eden Prairie, MN) with a steel hemisphere indenter (1 mm radius). Stepwise stress-relaxation tests (each step 5% nominal strain increment at a strain rate of 58% s^−1^ coupled with a 300 s relaxation period) were performed up to a strain of 20%. All tests were performed in a 1× PBS bath to maintain osmotic equilibrium. Relaxation period was set to ensure that stress equilibrium was achieved. The instantaneous modulus was calculated based on the largest compressive stress experienced during each strain step as previously described ^59^.

### AFM System and Data Collection

An AFM system (Keysight 5500, Agilent) was used for all experiments described. Needle-tip AFM cantilevers (NaugaNeedles) were modified from pyramidal tipped probes ^22^. The radius of curvature for the needle tip was approximately 25-100 nm and each needle was between 7-12 μm long with a precalibrated spring constant of 0.8 N/m.

### Probing HeLa CHMP4B-GFP with AFM

HeLa cells stably expressing CHMP4B-GFP were transferred to a tissue culture treated, low profile glass bottom dish (ibidi USA Inc., Madison, WI, USA) and incubated overnight before AFM testing to assure cells adhered sufficiently to the surface. Before testing, HeLa cells were stained with DAPI (1:1000) to visualize and target nuclear structures with the AFM needle-tip probe. To assure plasma and nuclear membrane puncture, a typical velocity of at least 10 μm/s and trigger force larger than 5 nN were needed. However, the deformable HeLa membrane required a higher velocity of up to 20 μm/s to assure double membrane puncture.

### Confocal Imaging of HeLa CHMP4B-GFP and AFM needle-tip probe

For visualization of the needle probe during cell membrane and nuclear envelope puncture, we treated the needle probe with 1 μg/mL fibronectin (Invitrogen) overnight at room temperature. The following day, probes were rinsed with DI water 3× and incubated with CellTracker Deep Red (C34565, Invitrogen) at room temperature for 30 minutes. Coating the needle in fibronectin followed by fluorescent labeling facilitated visualization of the needle probe. Probes were then mounted on the AFM for probing the HeLa CHMP4B-GFP cell line. The AFM was mounted on the confocal microscope with a custom adaptor on an optical table to minimize vibrational noise during AFM measurements. Thus, AFM was combined with confocal imaging for visualizing the cell/nuclear membrane puncture by the sharp AFM needle tip coated and fluorescently labeled in the far-red fluorescent channel. *Z*-stack images (512×512, 2 μm *z*-resolution) of a single HeLa CHMP4B-GFP cell were acquired before and during penetration (**Figure 2B**) with a laser scanning confocal microscope (Nikon A1, Nikon, USA) using a 40× dry objective.

### CHMP4B-GFP Recruitment After AFM Needle-Tip Penetration

Separately, we observed and imaged CHMP4B-GFP recruitment to the plasma membrane following AFM needle-tip puncture. We mounted the AFM system onto an inverted epi-fluorescence microscope (Nikon Ti-Eclipse) and used a 100× objective to target and image HeLa CHMP4B-GFP before puncture and after puncture at imaging intervals of 1 minute up to 10 minutes. The relative intensity at the puncture site was calculated by taking the ratio of the intensity signal at the wound site to the site 10 pixels above in the *xy*-plane that is undisrupted membrane (**Figure 2D**). For comparison of these relative intensity values, we used an AFM sphere-tip probe (5 μm borosilicate glass bead, Novascan) and a pre-calibrated spring constant of 0.07 N/m. The sphere-tip probe may indent a cell structure with the same setpoint force value as the needle-tip probe without inflicting a wound to the cell membrane. The ratio of the intensity at the contact site relative to the site 10 pixels above yields the intensity ratio for the sphere-tip probe.

### AFM Measurement and Data Analysis

Force-spectroscopy measurements for isolated cells and tissue slices were made in 1X PBS at room temperature. Measurements were completed within 1 hour of removal of culture media, and cell viability was confirmed in real-time with Calcein AM vital staining. The AFM probe center, containing the region with the needle, was aligned over a single cell structure in tissues or in cell culture populations. The AFM tip approached the sample and was stopped when deflection reached a very low setpoint (< 1nN), indicating the location where the needle tip meets the outer cell membrane, before increasing the setpoint force to initiate membrane puncture measurements. To determine cell membrane and nuclear membrane stiffness, a linear fit was performed over the force-relaxation segments in MATLAB (MathWorks) (**Figure 1C**). When pressing a cantilever to induce a large deformation of a soft cell structure, the effect of the substrate on the force curve is significant and contact area is variable so we did not apply a Hertz model to extract elasticity as is common practice for a spherical probe in biological samples. We, therefore, determined the stiffness of cell and nuclear membrane structures from the slope of the linear region of the force versus deformation curve ^60^. The same limitation that is poor estimation of contact area applied to measuring extracellular matrix stiffness with the needle probe, so these force-deformation curves were also fit to a linear model.

### AFM Stiffness Mapping of Isolated Chondrocytes

Primary chondrocytes were seeded onto a glass-bottom petri dish (ibidi USA Inc., Madison, Wisconsin, USA) coated with 1.5 µg/cm^2^ of fibronectin (Sigma-Aldrich, St. Louis, MO, USA). After incubation for 2 days to ensure strong attachment to plate, chondrocytes were ready for AFM testing and were immersed in sterile PBS after Calcein and DAPI staining, described above. The AFM system was mounted on a Nikon Eclipse Ti wide-field inverted microscope (Nikon Instruments) to simultaneously map stiffness and rapidly gather fluorescent images of testing areas. The needle-tip cantilever used in spectroscopy mapping experiments was pre-calibrated to 0.8 N/m by the thermal fluctuation method ^61^. After optically aligning the probe over an isolated chondrocyte, the AFM system was operated in force-volume mode to generate force spectroscopy stiffness maps, and each indentation was operated under these same parameters. An array of punctures and indentations were collected (32×32 indentations, 20×20 μm), wherein each indentation was operated to approach the sample at 15 μm/s and retract when the setpoint reached 5 nN. Each pixel of the resulting map represented one indentation puncture and the stiffness of each pixel was extracted by fitting the force-distance curve before relaxation to a linear model (**Figure 4**).

## Statistical Analysis

The apparent Young’s modulus for bulk cartilage measurements, and the calculated stiffnesses for AFM measurements of extracellular, cell membrane, and nuclear membrane structures are reported as mean ± standard error of the mean. Each experiment was analyzed by two-way ANOVA and Kenward-Roger test for pairwise comparisons in R 4.0.0 using the Estimated Marginal Means package. The statistical significance in each comparison was evaluated with p < 0.05 to denote significance.

## ACKNOWLEDGEMENTS

K. McCreery and X. Xu contributed equally to this work. We would like to thankfully acknowledge N. Emery, S. Nordstrom, and S. Schneider for advising appropriate models to use for statistical analysis. We would also like to thank J. Barthold for coordinating and executing animal dissections. This work was funded by NIH grants R01 AR063712 and AR071359, NSF CAREER grant 1349735, and with generous support from the W.M. Keck Foundation.

